# Systematic Comparative Analysis of Pathway Databases from a User’s Perspective

**DOI:** 10.1101/2025.03.26.645364

**Authors:** Moushumi Goswami, Divya Dosemane, Sravanthi Davuluri, Akhilesh Bajpai, Kshitish K Acharya

**Affiliations:** Institute of Bioinformatics and Applied Biotechnology (IBAB), Bengaluru – 560100, Karnataka, India; Research Scholar, Manipal Academy of Higher Education (MAHE), Manipal – 576104, Karnataka, India; Shodhaka Life Sciences Pvt. Ltd., Bengaluru – 560 100, Karnataka, India

## Abstract

Pathway Databases are essential particularly in the post-genomic era as they provide organized information about biological pathways and processes. However, there are too many pathway databases with varying features. These variations, stemming from multiple factors, create challenges for users in making objective selections among the databases. Hence, there is a need to systematically compare these databases from a user’s perspective. Indeed, there is a need to compile all such databases first. We first compiled pathway databases through a thorough literature search and shortlisted those suitable for human pathway studies. We designed a query set consisting of genes, pathways, and conditions and performed a quantitative comparison of the output from these shortlisted databases. Furthermore, to assess update frequency among the pathway databases, we curated potential novel genes involved in the MAPK pathway and compared them with the information available in pathway databases. The quantitative comparison of pathway databases showed that PathDIP yielded the highest number of pathways when queried with gene and condition names, followed by other databases. However, when queried for pathway names, Reactome yielded the highest number of pathways, followed by PathDIP, KEGG, and others. The results also support the notion that most scientists may have been making very arbitrary choice of pathway databases.

## INTRODUCTION

Pathway analysis is central to the understanding of the molecular basis of many cellular events, and a huge amount of information has been accumulated related to multiple pathways across species (Khatri *et al*., 2012; Kyriakis and Avruch, 2012; Plattner and Verkhratsky, 2018; Jia and von Wirén, 2020). In these contexts, it was essential to develop databases and computational tools (collectively referred to as resources hereafter) to store and analyze pathway information systematically. Hence, many bioinformatics databases, such as KEGG (Kanehisa and Goto, 2000) and Reactome (Joshi-Tope *et al*., 2005), have been developed.

Enrichment analysis of pathways for a set of genes of interest also plays a critical role in the current era of ‘omics’ (Paczkowska *et al*., 2020; Zhao and Rhee, 2023). Hence, pathway enrichment analysis tools such as DAVID (Dennis *et al*., 2003), have been created. However, there have been several such resources, and this seems to have created a few challenges for the users.

The pathway databases particularly vary in their contents and utilities (Labena *et al*., 2018). However, a reliable comparative account of these details is not available, even though some efforts have been made to compare the pathway databases - as elaborated below. Lack of clarity on the analogous resource features may be forcing the users to make very subjective selections of the databases as part of their studies. Such a possibility was indicated by our preliminary literature review, which showed a non-uniform usage of the available pathway resource. In addition, the reason for users’ preferences for any database among the resources does not seem to be clear in the literature; most researchers do not report the rationale of their choice among multiple options of pathway databases or software for analyzing their gene sets. Indeed, the choices made by scientists across available resources seem to be arbitrary. Our preliminary studies indicated that the current frequency of their usage does not appear to be proportional to their actual efficiencies.

Biologists may make a subjective selection among pathway databases mainly due to two reasons: a) there are enough articles reporting the use of certain pathway databases, and researchers tend to follow such precedence; b) even when they try to reason with the choice among the available options, they would soon realize that it is not practical, as researchers working on specific biological topics of their interest, to deviate their time and efforts into a painstaking comparative study of all pathway databases.

The situation seems to be common in the case of many other bioinformatics databases and software, referred together as resources, and a thorough periodic comparative analysis of each type of resource, particularly from a user’s point of view, is warranted (Bajpai *et al*., 2020; Acharya *et al*., 2024a, b, c). Quantitative comparison of relative output as a response to multiple types of common queries, i.e., the yield-coverages (Bajpai *et al*., 2020) can provide the required guidance for many scientists, for multiple types of biological research. Apart from general comparison of features, quantitative results from systematic analyses can provide a relative account of yield-coverages and, hence, a scientific basis for selecting among the options for specific analytical needs, and enhance the overall research efficiency. Elaborate efforts have been in this direction in recent years in the areas of different applications such as protein-protein interactions (e.g., Bajpai *et al*., 2020; Acharya et al, 2024b), literature search engines (Bajpai *et al*., 2011; Acharya et al., 2024c); alternative splicing tools (Acharya *et al*., 2024a), scRNA-sequencing and data analysis (Goswami *et al*., 2024) and other bioinformatics resources (Mathuria and Acharya, 2024).

It is not surprising that some scientists have paid attention to such a need in the case of pathway resources and attempted to compare the pathway databases. For example, in 2001, Wittig and De Beuckelaer reviewed general features of 27 tools that included pathway databases, genome databases, enzyme databases and so on (Wittig and De Beuckelaer 2001). They focused primarily on analysing the biochemical pathways provided by the pathway databases. The pathway databases KEGG, ExPASy, EcoCyc, PathDB, WIT and UM-BBD were compared for aspects such as content, data organization, information relevance, pathway representation methods, and available data analysis tools. An additional assessment of these databases was performed on their search options and query outputs. The study found KEGG to be the most user-friendly among the compared databases. About ten years after this report, another effort was made to compare five human metabolic pathway databases, including well-known ones like KEGG and MetaCyc. The comparison was done based on the number of genes, reactions and the EC numbers. They observed that the information provided by each is largely unique, with very little overlap. Several reasons were identified discrepancies – a) databases cover different aspects of metabolism, resulting in variations in their contents; b) databases might break down the same pathway into varying numbers of steps, creating mismatch; and c) There could be inconsistency in naming the metabolites that might create confusion. (Stobbe *et al*., 2011).

Later, another in-depth analysis was performed to compare the compounds, reactions and pathway contents of the KEGG and MetaCyc databases (Altman *et al*., 2013). This study showed that the number of compounds given by KEGG was significantly higher. However, there were more reactions and pathway content in MetaCyc than in KEGG. It concluded that the KEGG database is relatively incomplete in regard to the pathway information. In 2014, a command line tool was developed that automated the comparative analysis of metabolic pathways between organisms. Along with this, it could also list enzymes that were exclusive to the organisms (Porollo, 2014).

One of the most elaborative comparative assessments was done by Chowdhury and Sarkar (Chowdhury and Sarkar, 2015), who compared 19 signaling pathways across 24 pathway databases, along with a detailed study on two signaling pathways. The report presented a historical perspective of different pwDB developments and discussed pathway information as well as certain technical features such as pathway visualization formats and the availability of downloadable links. The study included comparisons via the use of two signaling pathways in the context of ease of retrieval of information when searched using the respective pathway names (Hedgehog and Notch signaling pathways). The authors identified unique features in various databases with the intention of helping users choose the appropriate database for their purpose.

However, the study does not help users to make informed decisions from several other perspectives. Another recent study (Marco-Ramell *et al*., 2018) reviewed the existing bioinformatics tools for metabolomics data analysis, metabolite databases, and disease-based analysis methods. The authors pointed out the incompleteness of the pathway databases.

Thus, there are multiple gaps in the current understanding of the utilities of various pathway databases, and a thorough comparative analysis is warranted. Indeed, surprisingly, even a systematic compilation of all pathway databases has not been reported yet. Hence, there is an urgent need to carefully list such databases and perform a more detailed quantitative comparative analysis such as the one carried out in the case of literature search engines (Bajpai, *et al*., 2011). If the resources are too many, one could consider a thorough comparison for at least one or two type(s) of application as done in the case of the protein-protein interaction databases (Bajpai *et al*., 2020). Such quantitative studies can also test if pathway databases have substantial differences in the outputs of at least in the context of certain types of applications. We have now taken a similarly detailed approach to first compile a list of pathway databases, compare their features, and systematically compare the yield and coverage of those relevant to human studies from a user’s perspective. We also test the hypothesis that the relative frequency of current usage of pathway databases is proportional to the relative output capacity of the databases.

## METHOD

### Database compilation, categorization and shortlisting

Multiple literature search engines (Bajpai, *et al*., 2011; Acharya et al, 2024a, b, c) were used to search and prepare a list of all pathway databases. PubMed, Google Scholar, and Google were the main search engines used with suitable query sets. Each database was used a few times to test its suitability for detailed analysis. Based on the main features and potential applications, the databases were categorized and many databases were found to be suitable for studying human pathway databases. These databases that implied our selection criteria, i.e., they are freely available online and give human pathways, were chosen for our study. Those that were commercial or whose URLs were not functioning were not included in the study.

### Query Selection

For the gene-based query, a combination of genes with ubiquitous as well as tissue-specific expression was selected. We also selected another set of genes that are associated with some of the well-known diseases (Bajpai, *et al*., 2020) and biological processes were selected, as described here. The mammalian gene expression databases [(Acharya *et al*., 2010) (MGEx-Tdb: http://resource2.ibab.ac.in/cgi-bin/MGEXdb/testisnew/Homepage.pl); (MGEx-Udb: http://resource2.ibab.ac.in/cgi-bin/MGEXdb/microarray/scoring/interface/Homepage.pl); (MGEx-Kdb: http://resource2.ibab.ac.in/cgi-bin/MGEXdb/kidneynew/Homepage.pl)] were analyzed to identify the genes based on their expression in tissues. Different disease-specific genes were collected from two online databases – eDGAR (Babbi *et al*., 2017) and UALCAN (Chandrashekar *et al*., 2017). Lastly a list of genes present in different biological processes were curated from literature. Diversity was ensured across all the genes in terms of the extent to which each gene is studied (i.e., the number of publications on each gene).

For the pathway-based query, diverse types of pathways related to signaling, metabolism, biochemical reactions, diseases, immunology, and DNA repair were first listed. A PubMed (https://pubmed.ncbi.nlm.nih.gov/) search was then performed to count the number of articles corresponding to each pathway. A set of pathways that included metabolic and various signaling pathways were selected to represent the diversity of biological relevance and the extent of studies done on each pathway.

Similarly, for the condition-based queries, literature was searched to select non-infectious diseases such as cancers, metabolic disorders, etc., and biological processes.

### Quantitative Comparison of Pathway Databases

The queries were searched in the databases, and the output was collected under the following headers: a) number of hits, b) number of pathways, c) number of relevant pathways, and d) name of the relevant pathways. An additional criterion was added for the pathway-based queries, where the number of genes was also collected for the relevant pathways.

The test of significance was performed to compare the means for the pathway-based query outputs between two databases at a time for top-ranking databases. As our data was non-parametric, the Mann-Whitney U test was performed.

### Quantitative Comparison of MAPK Signaling Pathway

We further decided to perform a detailed comparison for one of the pathways among the pathway queries. For this, a literature search was done to identify the pathway which is widely studied. This literature review led to the selection of the MAPK signaling pathway. For practical reasons, the top ranking databases shortlisted from the quantitative comparison were selected for the detailed comparison. These databases were then searched with the query ‘MAPK signalling pathway’. The relevant pathways were then chosen among the resulting hits, and genes present in the pathway(s) were collected. A non-redundant list was created for the number of genes per database.

### Qualitative Comparison of MAPK Signaling Pathway

Following the collection of the genes, we also wanted to explore the possibility that databases might miss out on some gene(s) while curating a pathway. Next, we categorized the MAPK pathway into phases by looking at the literature. The initiator phase would involve proteins (genes) that trigger the signal. These include growth factors such as EGF and PDGF along with their receptors – EGFR and PDGFR, respectively, in extracellular fluid, as well as the membrane proteins and ligands, and intracellular proteins (e.g. GRB2 and SOS) interacting with these membrane proteins. The intermediate phase would involve the proteins (genes) undergoing the signal cascade and carrying forward the signal. The intermediate phase mainly includes the kinases such as Ras, MAPK, and ERK, which are central to the MAPK pathway. Finally, the effector phase would involve proteins (genes) that bring about the required changes. This phase mostly includes transcription factors such as c-myc and c-fos, which in turn interact with DNA to elicit the final response. The genes collected from the different databases previously were now distributed under these phases based on their category definition. Literature was searched extensively during this task.

### DAVID-like analysis using PathDIP

Functional enrichment of gene sets obtained via genomic, transcriptomic, or proteomic approaches may provide different results depending on the pathway database used. To find the potential impact of leaving out some of the databases that performed well in the current study, we compared the results of the functional enrichment routinely performed by scientists using different databases. We tried to reproduce the results of some of the earlier, successful transcriptomic analyses first and repeated the analyses with the left-out database, PathDIP. Recent articles reporting NGS-based transcriptomics with functional analysis of three cancers that meet the following criteria were selected.

The formula used by the standard pathway analysis tool, DAVID (https://david.ncifcrf.gov/), was applied to the relevant commonly used databases KEGG and Reactome. We first tried to replicate the results in the studies, which was successful. Once we were certain of the formula, it was then applied to the PathDIP database outputs for the same lists. The results were compared.

## RESULTS

A comprehensive list of 185 pathway databases has been created and categorized in multiple ways based on their features and the type of information these pathway databases provide (https://startbioinfo.org/cgi-bin/resources.pl?tn=Pathways). The usage frequency of the databases was assessed using a standard procedure (https://startbioinfo.org/description.html). The results show that KEGG is the most commonly used database (91%) for human-related pathways and information. Among other databases (9%), Reactome is the most widely used, followed by PharmGKB (20%), Panther (19%), and WikiPathways (5%), while most others are hardly being used.

An initial list of 45 online pathway databases (pathway databases) that were free to use were selected for the comparative analysis. However, as indicated in the workflow (Figure 1), some of these databases were excluded as they did not comply with our selection criteria. A couple of databases were very specific to a particular topic. Some databases had also stopped functioning during the course of our study. The lack of functionality in such cases was confirmed by repeated attempts, across several months, before eliminating them from further consideration. Eventually, only 14 pathway databases could be compared systematically according to the plan.

**Figure 1.**
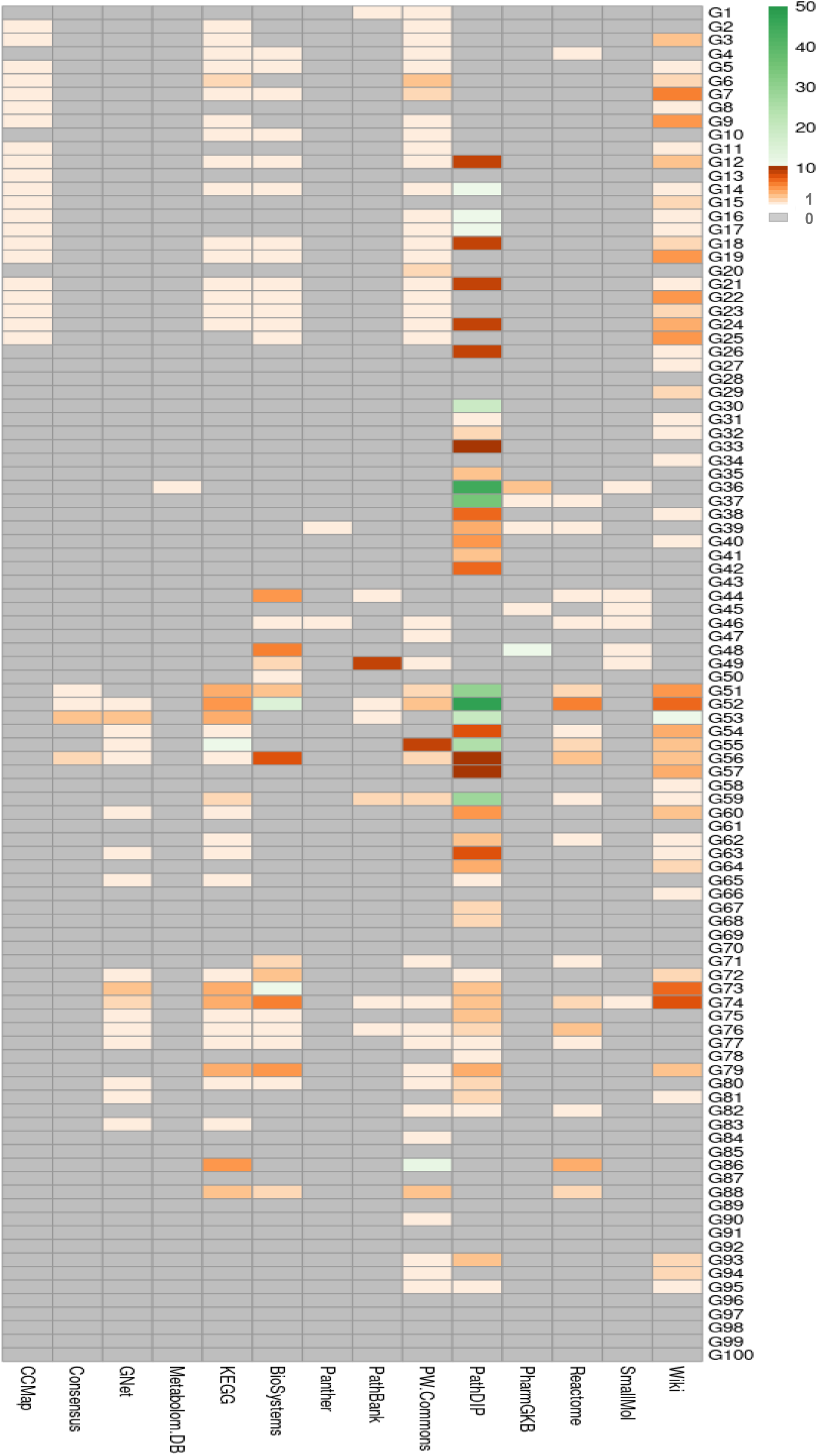
The relative number of relevant pathways obtained in response to individual gene queries (100 genes).

For the gene-based query, less than 50% of the gene names were covered across the DBs. Around 40% covered by WikiPathways, PathDIP, Pathway Commons, 20-30% were covered by KEGG, NCBI BioSystems and Cancer Cell Map. Other databases covered less than 20% of genes. Although PathDIP covered a total of 48 genes, it gave the highest number of pathways for the genes, which is around 446. In fact, this pwDB stands as an outlier when compared to the output of the other databases, as the next highest number of pathways given by any database is 126 (CPDB), which is less than half of the pathways covered by PathDIP. Thus, PathDIP stands out from the other databases when it comes to gene-based queries. It should be noted that the PathDIP database provided literature-curated information and covered experimentally proved and predicted protein-protein interactions. For the pathway-based query, around 50-60% of the pathway names were found WikiPathways, CPDB and Reactome. Around 40% of pathway names were found in PathDIP. The rest of the databases gave less than 40% of the pathway names. Reactome covered only half of the total pathway queries. However, it gave the highest number of pathway outputs when compared to the rest of the databases. It actually stands out by giving more than 800 pathways for the queries. The next highest pathways were given by CPDB, which is less than 25% of the result given by Reactome. For the condition-based query, 70-85% of the condition names were found in WikiPathways, followed by Cancer Cell Map and KEGG. 30-60% of the condition names were found in other databases. However, Panther did not respond to any condition names used in the current set of queries. Although pathDIP covered only 50% of the condition names, the highest number of pathways were obtained. This database was followed by ‘cancer cell map’, HMDB, KEGG, and WikiPathways.

### Comparisons with gene queries

The number of relevant pathways retrieved per gene across pathway databases, as well as the number of pathway databases responding across genes, varied substantially (Figure 1).

Surprisingly, querying with none of the genes resulted in relevant pathways from all of the 14 pathway databases. Indeed, a maximum of nine pathway databases provided such an expected positive result for a queried gene, and this was the case for the genes ZAP70, RB1, and AURKA. Three other genes (XPC, BRCA1, and FGFR3) received positive results from seven pathway databases each, while six genes (RPL24, RPL27, RPS24, WFS1, CHECK2, and AMACR) received such results from six pathway databases. While two to five pathway databases responded positively for 11 to 16 genes, only one pwDB provided positive results in the case of 19 genes. PathDIP gave results for eight of these genes, WikiPathways for five, Pathway Commons for four, and Cancer Cell Map for one gene.

PathDIP was the top performer that provided the highest number of relevant pathways (446) in response to 48 of the 100 genes queried. WikiPathways was the next best, responding to 49 genes, resulting in 126 relevant pathways. NCBI BioSystems, Pathway Commons, and KEGG responded positively to 30 to 45 query genes and yielded 70 to 90 pathways each. The next level of performers were Reactome, Cancer Cell Map, GenomeNet, PharmGKB, and PathBank, with 15 to 35 relevant pathways from each DB, while responding to 5 to 21 genes. Human Metabolome Pathway Database, Panther, ConsensusPathDB, and The Small Molecule Pathway Database, revealed less than 10 relevant pathways less than 10, while responding to 1 to 7 genes.

There were some genes (e.g., SLC12A1, RB1, MMP7, PCNA, etc.) to which multiple pathway databases provided a high number of relevant hits, and there were some pathway databases that responded to the highest number of genes, there was no consistent trend. Exceptions included genes for which only the good performer, such as PathDIP (for LDHC), or a poor performer, such as PantherPanther (for RPL9) provided any result. Thus, to ensure complete coverage, the user may have to use multiple, if not all, of the 14 pathway databases.

### Comparisons with condition-queries

The number of the relevant pathways retrieved per condition used as a query and the number of genes revealed across hits varied substantially across pathway databases (Figure 2). Similarly, the number of pathway databases responding across qPWs was also diverse.

**Figure 2.**
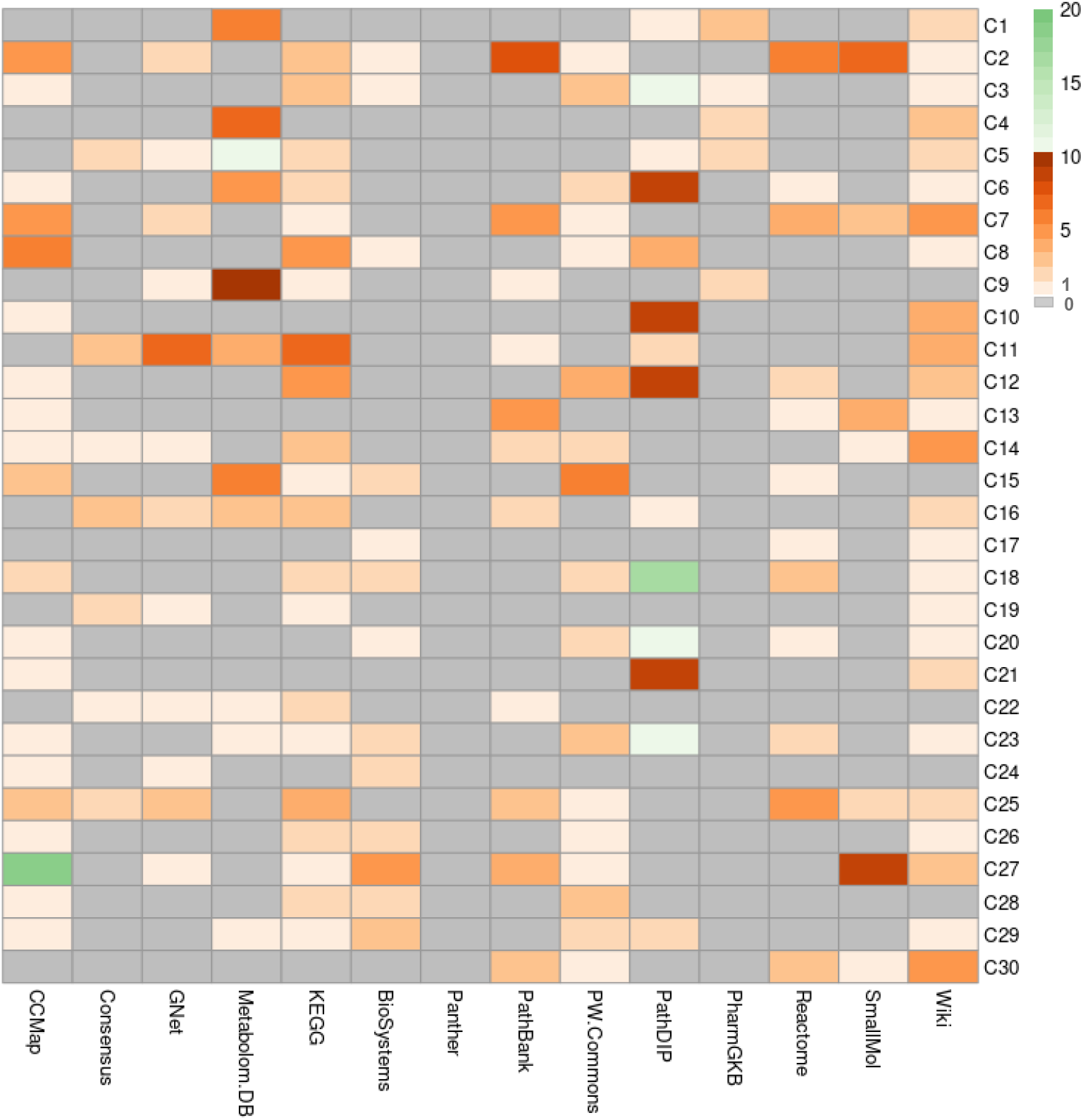
The relative number of relevant pathways obtained in response to individual conditions as queries (30 conditions).

PathDIP again performed well with the highest number of relevant pathways (94). It responded positively to 15 of the 30 condition queries. Reactome also responded to a similar number (13) of queries but resulted in 27 relevant pathways. Cancer Cell Map and Human Metabolome Pathway databases revealed 55 and 54 relevant pathways in response to 21 and 11 conditions, respectively. Among others, WikiPathways and KEGG gave positive results for 70-80% of the queries; however, each of them output only 50 relevant pathways. Other pathway databases yielded the relevant pathways in a range between 10 and 34, and these included PathBank (34) and Pathway Commons (33).

It is difficult to recommend one database over the other due to the variations in the results. For example, PathDIP, the top performer, does not respond well when queried with Meiosis. On the contrary, CCMap, the Small Molecule database, and many other databases provided a decent number of positive hits. Thus, as seen in the case of genes, it may be better to use a combination of multiple databases even when querying with conditions.

### Comparisons with pathway queries

The number of relevant pathways retrieved per PathWay name used as a query (qPW), as well as the number of genes revealed across hits, varied substantially across pathway databases (Figure 3). Similarly, the number of pathway databases responding across qPWs also varied. In terms of the number of relevant pathways among the results, Reactome emerged as the top performer with an exceptionally higher (884) number of pathways while responding to 25 of the 50 qPWs. Even though the high number was an outlier number of 750 pathways in response to one pathway (protein kinase pathway) only, the total number across other pathways was still higher (234) than the output from other pathway databases. A total of 1254 genes were covered among the relevant pathways revealed by Reactome in its result pages. A maximum number of genes (1910) were covered among the relevant pathways revealed by PathDIP even though it provided only 95 total such pathways in response to 21 qPWs. The gene count was also higher (1477) in KEGG results among its relevant pathway hits (13) despite the response being seen for 14 qPWs only. In contrast to this was ConsensusPathDB, with only 447 genes across 178 relevant pathway hits in response to 27 qPWs.

**Figure 3.**
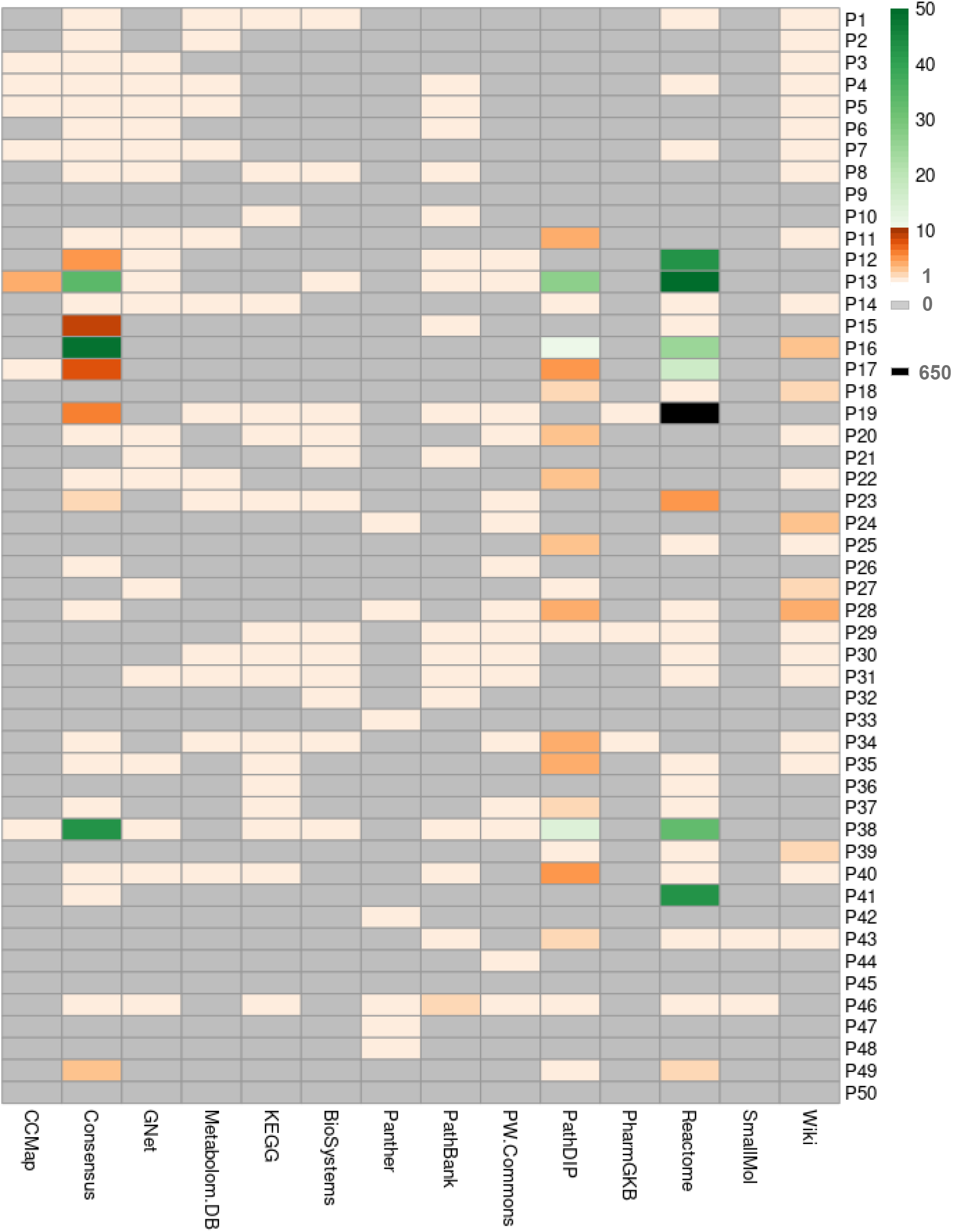
The relative number of relevant pathways obtained in response to individual pathway names as queries (50 pathways).

WikiPathways also revealed 1474 genes among its relevant hits (38 pathways) while responding to 28 qPWs, and GenomeNet results covered 1012 genes among 18 hits for 19 qPWs. The PharmGKB, Human Metabolome, and Small Molecule Pathway databases covered 61, 41, and 3 genes, respectively. Of these three, only the Human Metabolome database responded to 14 qPWs, while the other two responded to only two or three qPWs.

Overall, there was no correlation between the number of qPWs a database responded to, with either the number of relevant pathway hits or the genes covered among the hits. The performance of the 14 pathway databases performed varied depending on the parameters assessed. As in the case of gene queries, there were some qPWs (e.g., p53, MAPK, and Notch) to which multiple pathway databases provided a high number of relevant hits, and there were pathway databases that responded to a higher number of qPWs. But there was no such consistent trend. Exceptions included qPWs for which only a good performer, such as Reactome (for pathway PD-1) provided a good number of relevant hits. Panther, which responded to only six qPWs, responded exclusively to two qPWs (p16 and GABA). Thus, to ensure comprehensive coverage, the user may have to use multiple pathway databases when querying with pathway names as well.

Across the three types of queries, PathDIP yielded the highest quantum of information consistently, even though Reactome and ConsensusDB yielded more positive hits than PathDIP in response to pathway names. A consistent good performance across query types was evident in the case of PathDIP followed by Wikipathways, while Reactome performed better only with pathway name and condition-based queries. It should be noted that the total number of genes covered among the pathway hits, in response to pathway-names as queries, was highest again in case of PathDIP. The results suggest an advantage in using multiple databases for any of the three query types while suggesting one or a few better performers.

Since the number of genes covered by each pathway revealed is critical, in response to any query, we analyzed responses to qPWs in detail. The goal was to assess exclusive coverage of the genes across pathway databases. Results of a subset of ten pathway names were compared in detail, and the following observations (figure 4) were made:

**Figure 4.**
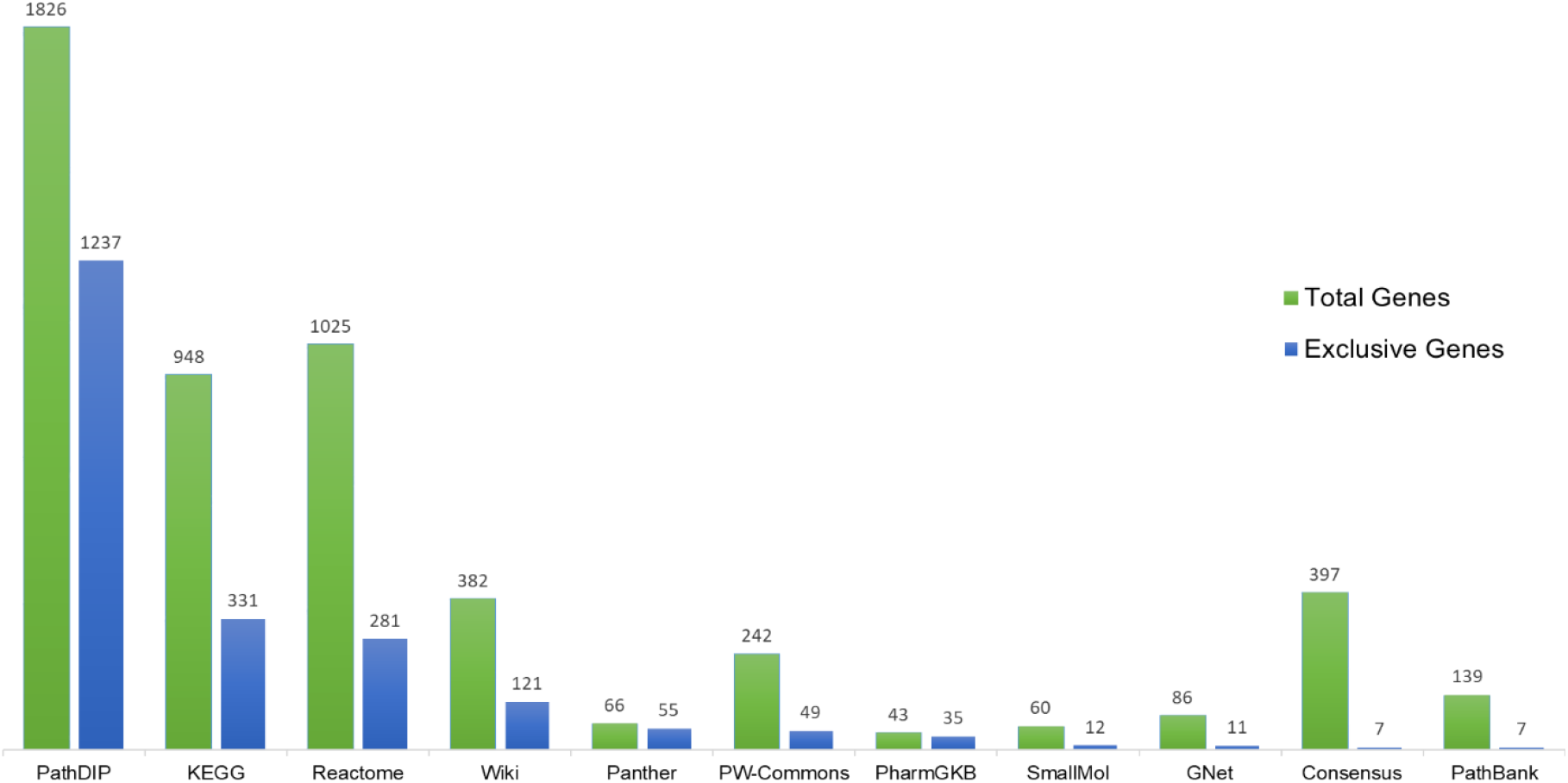
A comparison of the total and exclusive number of genes revealed across the databases in response to querying with ten pathway names.

PathDIP again provided the highest number of genes (1826) and contributed the highest number of exclusive genes (1237). The next best contributors were Reactome (1025) and KEGG (948). The total number of genes, as well as exclusive contributions, were less in the case of other databases.

### Quantitative Comparison using MAPK signaling

To further compare the gene coverage across databases and literature, a more detailed study was conducted using the MAPK pathway as a case study. This laborious quantitative analysis of one pathway was restricted to six better-performing databases. Reactome gives the highest number of total genes (657) for the MAPK pathway, followed by PathDIP (590). While this trend partially contradicts the observations made with 50 qPWs, where PathDIP provided the best results, the other pathway databases seem to perform in accordance with earlier observations with a lower number of genes. A lower total number of genes was revealed by KEGG (294), Pathway Commons (252), ConsensusPathDB (65), and Panther (30).

Compared to the rest of the pathway databases, Reactome provided the highest number (303) of exclusive genes compared with PathDIP. Pairwise comparisons revealed interesting results. For example, a total of 327 genes covered were common between Reactome and PathDIP. Reactome covered 330 genes in the MAPK pathway that PathDIP did not cover, whereas PathDIP covered 263 such genes that Reactome did not cover.

A qualitative comparison of the MAPK pathway was also performed across pathway databases. After comparing databases and information from the literature, three main phases of the pathway were identified: the initiator phase, where growth factors interact with membrane-bound receptors and cytoplasmic proteins interact with such receptors; the intermediate phase, with protein interactions in the cytoplasm; and the effector phase, involving gene activation.

Information from different databases was compared, and KEGG and PathDIP showed better coverage of genes across all three phases considered. The latter simply reflected KEGG in the case of the MAPK pathway. Hence, the results were identical. Pathway commons and Reactome also performed similarly with a slightly lesser number of genes.

The rate of updating the information in the databases and potential loss of information available in the literature are also essential aspects that concern users. Hence, we undertook a study to collect information from the literature about protein interactions involved in the MAPK pathway and compared the output with different databases.

Using the already existing list of MAPK genes from the better-performing databases, a union list of MAPK genes was made with a total of 900+ genes. Of them, 16 genes from this list were randomly selected. Research articles related to each of these genes were then searched. Articles were carefully screened, and information about the interaction of the proteins of concern with other molecules, preferably proteins, was gathered. A total of 279 molecules interacting with 16 query proteins were identified and compared with existing information in the database. Among these interactions, 201 were not covered by any database. The interactions listed included receptors, inhibitors (molecules and synthetic compounds), and post-transcriptional and post-translational modifications.

### A comparison of pathway enrichment analysis using the outputs from different pathway databases

Four cancer data-sets were identified for the study and the corresponding lists of differentially expressed genes were collected. Each of these gene-lists was then submitted to DAVID software as well as PathDIP, to obtain the list of represented pathways. For the DAVID-like enrichment analysis using the PathDIP database, an online tool was used to perform the calculation of the p-value for the modified Fisher Exact Test (https://www.socscistatistics.com/tests/fisher/default2.aspx). Another tool was used to derive the FDR and Benjamini and Hochberg values (https://tools.carbocation.com/FDR).

DAVID uses the linear step-up method of Benjamini and Hochberg (Benjamini and Hochberg, 1995) to obtain adjusted p-values. The FDR calculations in DAVID use adaptive linear step-up adjusted p-values for approximate control of the false discovery rate, as discussed in Benjamini and Hochberg (Benjamini and Hochberg, 2000). The lowest slope method for estimation of the number true NULL hypothesis is used.

The results showed that PathDIP analysis yielded a much higher total number of pathways for the same sets than the DAVID results. Also, the overall significant pathways obtained with PathDIP were higher than those obtained with other databases used in DAVID. When checked for significant exclusive pathways, Reactome gave the highest number, followed by PathDIP. However, when the same sets were submitted in PathDIP, the number of significant exclusive pathways was highest.

## DISCUSSION

The current study forms the most detailed comparative analysis of pathway databases (pathway databases) to date, which is also the first to go beyond simple feature comparisons to perform a quantitative comparison of yield-coverage of pathway databases with multiple query types. Even though, for practical reasons, such quantitative comparison was made on key databases applicable to human studies, substantial novel observations were made by extensively using three categories of queries, viz., genes, pathways, and conditions. The queries for each type were not only reasonable in size, but represented a good diversity of multiple perspectives including human biology and user-interests. The MAPK signaling pathway has also been used as a case for a more detailed assessment of the completeness and organization of information across pathway databases. The results demonstrate an advantage of specific pathway databases for specific types of applications and illustrate the need to use multiple databases to ensure higher coverage of pathways and genes.

One of the reasons for the lack of such detailed, quantitative comparative studies in the past is the laborious nature of the work required. Querying the databases individually is a painstaking task in itself as witnessed in the current study. Apart from making 4680 individual queries, the output from each query had to be analyzed carefully. In addition, most of the databases differed in terms of information retrieval. For instance, 30 pathway hits would be obtained in Reactome when queried with the phrase ‘Insulin signaling’, and ‘Signaling by insulin’ was chosen as relevant. Another example can be taken of the ‘TWEAK (*Tumor necrosis factor-like weak inducer of apoptosis*) pathway’ name, which when queried, only NCBI BioSystems and PathBank provided relevant results. Others mostly provided interaction pathways of cytokines, TNFs, or no result at all. Choosing a relevant pathway was challenging in cases like this, as TWEAK is a cytokine that uses TNF receptors for signaling. As a matter of fact, if this pathway would appear while querying the cytokine pathway, it would be considered relevant. However, as we are specifically searching for the ‘TWEAK pathway’, other cytokine-based or TNF-based pathways were not considered relevant. Similarly, when the ‘MET (*Mesenchymal Epithelial Transition*) pathway’ name was queried, except for Reactome and WikiPathways, which provided ‘*Signaling by MET’*, all the other DBs either provided results for ‘*methionine metabolism*’ or no results at all. Such results were due to the standard three-letter abbreviation of the amino acid methionine, which MET/met. Once the pathway was queried, all the pathway names provided by the DBs were listed, followed by filtering out the relevant ones among the rest. This filtering is even more difficult when it comes to conditions; in fact, one of the hardest times selecting relevant pathways was for a condition. This difficulty is caused by the involvement of more than one pathway. For instance, most signaling pathways are involved and dysregulated in any type of cancer. When queried with ‘Gastric Cancer’, only some DBs provided straightforward results like ‘*Gastric cancer network*’ (ConsensusPathDB, GenomeNet, PathDIP) or ‘*Gastric Cancer*’ (KEGG and WikiPathways). Databases such as the Human Metabolome Database and PharmGKB provided pathways like ‘*Methotrexate action pathway*’ and ‘*Doxorubicin metabolism pathway*’, among others, which demonstrate the pathway through the mechanism of action of a drug or type of treatment when queried a condition. This trend was observed for most of the conditions in our query list. When queried with a biological process under the condition query, the results were equally complex in identifying which pathways are involved in them. For instance, when queried with spermatogenesis, Reactome provides pathways like ‘*Regulation of endogenous retroelements by Piwi-interacting RNAs (piRNAs)*’ which is a very broad pathway, PathBank provides pathways like ‘*Intracellular Signalling Through FSH Receptor and Follicle Stimulating Hormone*’, and KEGG provides pathways like ‘*Oocyte meiosis*’, ‘*Progesterone-mediated oocyte maturation*’, none of which are pathways associated with spermatogenesis. Other DBs do not even give any result output for spermatogenesis. All these pathways were considered to be irrelevant for spermatogenesis.

Sometimes, when a pathway was queried, it responded with the process it was involved in; such results were considered relevant. An instance of this is while querying with the ‘Kisspeptin pathway’, which is a type of neuroendocrine pathway that aids in the regulation and release of GnRH. The DBs (KEGG and Reactome) that responded with a result for this pathway provided the ‘*GnRH secretion*’, and it was considered relevant, as it demonstrates the process the pathway is involved in. Similarly, for cAMP pathway, the pathways like ‘*Activation of cAMP-dependent protein kinase*’ (PathBank, PathDIP and Pathway Commons) was considered to be relevant. Other DBs (GenomeNet, KEGG, PathDIP, and NCBI BioSystems) responded with the ‘*cAMP signaling pathway*’.

These variations across pathway databases can be a problem for users. Another type of variation was observed while querying with the name of the pathway, which may be stored as different phrases in a particular DB. For instance, the phrase “MAPK signalling pathway” applicable in KEGG seems to be equivalent to “MAPK family signalling cascades” in Reactome, where the MAPK pathway has been broken into various sub-pathways like MAP2K and MAPK activation, negative regulation of MAPK, and so on. However, KEGG just responds with one pathway.

There are also alternatives of a particular pathway that were observed in various databases such as MAPK signaling pathways can also be found under the names of ‘*ERK1/2 signaling pathways’* (Guo *et al*., 2020; Braicu *et al*., 2019) or ‘*P38 MAPK pathway’* (Braicu *et al*., 2019; Dodeller & Schulze-Koops, 2006) in some of the databases like PathDIP, Pathway Commons, WikiPathways, etc. Similarly, the mTOR pathway can also be found under names of ‘*PI3K-AKT-MTOR pathway’* in PathDIP, and ‘*Focal adhesion p13-akt-mtor signaling’* in Reactome. The reason for this is due to the connectivity of mTOR with the PI3K-AKT signalling pathway (Glaviano *et* al., 2023; Paplomata & O’Regan, 2014). However, TCA cycle is one of the pathway query names, where all the DBs responded with the pathway with the same name as the query.

Still, we found two exceptions of the Human Metabolome Database and Reactome, where the pathway is curated under the name of ‘*Citric acid cycle’* (Martínez-Reyes *et al*., 2020; Akram, 2014). While querying the pathway Double-strand break repair in KEGG, it provided ‘*Homologous recombination (HR)’* and ‘*Non-homologous end joining (NHEJ)’* in response.

However, Reactome responded with the same name as queried, along with *‘DNA Damage Repair’*. It is known that the double-strand break repair (Tan *et al*., 2023) mechanism can occur via *HR* (Krejci *et al*., 2012; Elbakry & Löbrich, 2021) or *NHEJ* (Taleei & Nikjoo, 2013; Chang *et al*., 2017), so, either of them can be considered relevant for this specific query.

We also observed the spelling of signalling differs, some databases have used ‘signalling’ and some have used ‘signaling’. For instance, when the word ‘signaling’ is used in Reactome, it just considers the prefix (pathway name) attached to it to search for the pathways. This also changes the initial number of hits obtained after the querying. When given the word ‘signaling,’ it considers the whole query as a phrase to search the pathways. Although there is a change in the initial hit, it does not reflect upon the final relevant pathways. While such an observation may be relevant for current discussion, such aspects do not form a major concern when it comes to the relevance of pathways for a particular query.

Detailed analysis was also needed when identifying relevant pathways using gene names as a query. The gene did not play a central role in the pathway, connected to most of the important pathway components in the primary axis, in all cases. For example, when the AURKA gene is queried in Pathway Commons, two relevant pathways provided by the databases were ‘*AURKA Activation by TPX2’ and ‘Interaction between PHLDA1 and AURKA’*. But the two other pathways, viz., *‘TP53 Regulates Transcription of Genes Involved in G2 Cell Cycle Arrest’* and ‘*Generic Transcription Pathway’* were irrelevant, as AURKA was not even in the secondary axis of these pathways. However, we understand that such relevancy calls can be subjective as emphasized below.

In many cases, we needed to identify the “relevant” ones among the others, which we did by verifying with information from the literature. For example, even though many DBs responded to ‘CHEK2’ with ‘*ATM signaling* pathway’, some DBs yielded multiple other responses that required some amount of literature review or review of the pathways themselves before deciding the relevance. Such pathways included *cellular senescence* in KEGG *and DNA damage-induced protein phosphorylation, regulation of DNA damage checkpoint* in Cancer Cell Map, *Cell cycle* in GenomeNet, *regulation of cell cycle progression by plk3, role of brca1 brca2 and atr in cancer susceptibility* in Pathway Commons.

Understanding the variety of functionalities of pathway databases was also essential. For example, while all pathway databases could be queried with official gene symbols, KEGG used its abbreviations or symbols when representing genes in the output pathways, which made it difficult to track and count genes among pathways.

The final calls on relevancy can be made differently by different researchers. We had to make some rules for the sake of uniformity. For example, for Alzheimer’s condition, Reactome showed in one relevant pathway, viz, *Deregulated CDK5 triggers multiple neurodegenerative pathways in Alzheimer’s disease models* (Liu *et al*., 2016). We decided that ‘*Defective Base Excision Repair Associated with OGG1’* (Wang *et al*., 2025) and ‘Defective *OGG1 Substrate Processing’* were irrelevant. Even though there were a good number of articles linking oxidative damage of DNA with neurodegeneration, and several implicating OGG1 (e.g., Abolhassani *et al*., 2017), we did not consider the two pathways involving OGG1 to be relevant to the Alzheimer’s condition as they are primarily DNA repair (Base-excision repair) pathways (Wang *et al*., 2018; Jacob *et al*., 2013). The priority in this study was to identify what is definitely relevant and not to take a definitive call on what is irrelevant.

It is important to note that the relative usage frequency of pathway databases by the scientific community contradicted the yield-coverage results, particularly with respect to KEGG, which seems to be used most frequently. In contrast, PathDIP, which appears to be used sparingly, provided the best yield-coverage with at least two types of queries. This latter situation may also be due to the follow-up work required with PathDIP to compile results from multiple sources when using this database or simply to lack awareness. ConsensusPathDB and Pathway Commons also performed unexpectedly well. However, the usage frequency did not contradict the currently observed advantages of KEGG, Reactome, and Panther, which performed relatively well compared to many other pathway databases.

The current study also involved preparing the most comprehensive list of pathway databases (pathway databases) with 192 pathway databases listed at startbioinfo.org (https://startbioinfo.org/cgi-bin/simpleresources.pl?tn=Pathways). Shortlisting pathway databases for final comparison required careful analysis. Among the functional pathway databases relevant to human studies, those specific to a certain topic were excluded. Examples of such databases include Modomics (specific to RNA metabolism and synthesis), RepaireTOIRE (DNA repair pathways), and T1D database (type 1 diabetes). Databases, such as SignaLink and XTalkDB, with limited gene information and mostly focused on the crosstalks observed in the signaling pathways, were also excluded. In addition, pathway databases were selected for the final comparison only if they provided results for all three query types. Exclusions in this round included Molecule pages and InnateDB. MetaCyc required a subscription after a certain number of uses and, hence, was also excluded from our final study.

While best efforts have been made in preparing this list, we noticed continued development of new resources throughout the prolonged study period. For example, when the manuscript was being finalized, new databases such as SPathDB (Li *et al*., 2025), Enteropathway (Shiroma *et al*., 2024). WikiPathways has been recently updated as well (Agrawal *et al*., 2024). There also have been recent efforts to explore the utility of machine learning approaches in enhancing pathway-associated information (Robben *et al*. 2023; Neelakandan and Rajanikant, 2024). While several databases were found to be non-functional in the beginning of the study, it is also interesting to note that many functional ones stopped functioning during the period of work – a phenomenon we noticed while comparing other resources (e.g., Mathuria and Acharya, 2024). Examples include the BIOPYDB database that stopped working during our study (around 2021) and the NCBI BioSystem, which was retired in April 2022.

The constant changes in the available and functional pathway databases only reiterate the need for a periodic listing of pathway databases, their cataloging from a user perspective, and in-depth detailed comparisons for different types of applications. The current study compares online, publicly available databases for analyzing human pathways. Systematic studies are also needed for other species, and it is best to include commercial resources in such studies.

A general review or comparison of features alone will not help biologists much. Earlier pwDB comparisons did not cover enough queries or databases. Among such earlier mentioned studies, only one compared a high number of pathway databases. (Chowdhury and Sarkar, 2015). This study discusses the history of these databases, the pathway information they contain, and the built-in tools they offer. To further evaluate the strengths and weaknesses of these features in a practical setting, a case study on the Hedgehog and Notch signalling pathways was included.

While this study was a comprehensive one, several limitations can be highlighted in this study as well. First of all, most of the databases used in this study are either non-functional. Second, it compared only signalling pathways. Thus, many other aspects of pathway databases were not studied. Lastly, the comparison only looked at the presence or absence of a certain pathway in the database.

The current results may fill a critical void in pathway analysis. The results of the current study can help users to make several objective selections. For example, if only one database has to be used for gathering multiple specific pathways when querying with a pathway name, Reactome would be a good choice. PathDIP – a secondary database that sources information from more than 20 other databases - including KEGG and Reactome, can be a similar choice when querying with gene or condition queries – even though it requires a follow-up compilation. However, we recommend a combination of databases to minimize the loss of information due to multiple observations made in the current study.

The lack of responses to many queries is a matter of concern across pathway databases. For the gene-based search, none of the pathway databases elicit responses for all genes. Fourteen pathway databases did not respond to 15% of the gene names, whereas nine DBs responded to a maximum of 3% of the gene names. The highest number of genes covered by at least one DB was only 19%. For the pathway-based search, 8% of the pathway names did not elicit a response from any of the 14 databases, whereas nine pathway databases responded to a maximum of 4% of the pathway names found. The highest number of pathways covered by at least three DBs was 16%. For the condition-based search, although there was no such condition name that was not found by any DB, however, only 16% of these were found in at least three DBs. Around 6% of the condition names were found in a maximum of nine DBs. The highest number of conditions covered by at least six DBs was almost 23%.

The comparative studies would have been incomplete without a comparison of the organization of information, particularly in the context of relative abundance of information at different phases of at least one signaling pathway, and a comparison with the information in the literature. Hence, MAPK signalling pathway was compared in such details across the top performing database to analyze the differences in the gene coverage. KEGG, PathDIP, Pathway Commons, and Reactome showed better coverage of genes across all three phases considered compared to the remaining pathway databases. However, it is important to note that a substantial number of molecules involved in the pathways, which can only be detected via a thorough literature search, can be missed using any database. Hence, a researcher aiming to minimize loss of information would face a huge challenge of biocuration in addition to a combinatorial use of pathway databases. The observations also indicate a need to enhance efforts to update the existing pathway databases.

The DAVID database is very commonly used for pathway enrichment for a set of genes of interest derived via multiple omics studies. However, it covers just five pathway databases. On the other hand, PathDIP covers more than 20 databases and enables users to perform their own pathway enrichment analysis. In this context, comparing the pathway enrichment results of DAVID with those from PathDIP becomes significant.

The vastness and ever-changing nature of pwDB availability, as well as their features, make comparative studies extremely difficult. However, the current results demonstrate the necessity of conducting such studies periodically. Indeed, the need and benefit of such quantitative comparisons from a user’s perspective have been highlighted for many other types of bioinformatic resources (Acharya et al. 2024a, b, c; Mathuria and Acharya 2024; Bajpai et al., 2011).

## ACKNOWLEDGEMENT

Research projects at IBAB are supported by the Department of IT, BT, and S&T, Government of Karnataka, India. Ms. Vasumathi Manivelan, Department of Biotechnology, M S Ramaiah Institute of Technology, Bengaluru, Karnataka, India, generated heatmaps.

